# Less is worse than none: ineffective adaptive foraging can destabilise food webs

**DOI:** 10.1101/2021.11.28.470273

**Authors:** Hsi-Cheng Ho, Samraat Pawar, Jason M. Tylianakis

## Abstract

1. Consumers can potentially adjust their diet in response to changing resource abundances, thereby achieving better foraging payoffs. Although previous work has explored how such adaptive foraging scales up to determine the structure and dynamics of food webs, consumers may not be able to perform perfect diet adjustment due to sensory or cognitive limitations. Whether the effectiveness of consumers’ diet adjustment alters food-web consequences remains unclear.
2. Here, we study how adaptive foraging, specifically the effectiveness (i.e. rate) with which consumers adjust their diet, influences the structure, dynamics, and overall species persistence in synthetic food webs.
3. We model metabolically-constrained optimal foraging as the mechanistic basis of adaptive diet adjustment and ensuing population dynamics within food webs. We compare food-web dynamical outcomes among simulations sharing initial states but differing in the effectiveness of diet adjustment.
4. We show that adaptive diet adjustment generally makes food-web structure resilient to species loss. Effective diet adjustment that maintains optimal foraging in the face of changing resource abundances facilitates species persistence in the community, particularly reducing the extinction of top consumers. However, a greater proportion of intermediate consumers goes extinct as optimal foraging becomes less-effective and, unexpectedly, slow diet adjustment leads to higher extinction rates than no diet adjustment at all. Therefore, food-web responses cannot be predicted from species’ responses in isolation, as even less-effective adaptive foraging benefits individual species (better than non-adaptive) but can harm species’ persistence in the food web as a whole (worse than non-adaptive).
5. Whether adaptive foraging helps or harms species coexistence has been contradictory in literature Our finding that it can stabilise or destabilise the food web depending on how effectively it is performed help reconcile this conflict. Inspired by our simulations, we deduce that there may exist a positive association between consumers’ body size and adaptive-foraging effectiveness in the real world. We also infer that such effectiveness may be higher when consumers cognise complete information about their resources, or when trophic interactions are driven more by general traits than by specific trait-matching. We thereby suggest testable hypotheses on species persistence and food-web structure for future research, in both theoretical and empirical systems.

## Introduction

The structure and dynamics of food webs emerge from foraging interactions among organisms. It is therefore necessary to consider mechanisms that govern consumers’ diet choices to better understand the emergence of these food-web properties (Beckerman et al. 2006; Berlow et al. 2008; Stouffer 2010; Ho et al. 2019). It has been shown that real-world food webs are not static networks, but rather change over time and space as species “rewire” their trophic interactions (Blanchard 2015; Tylianakis & Morris 2017; Bartley et al. 2019). One predominant behavioural mechanism driving such rewiring is adaptive foraging (Uchida et al. 2007; Valdovinos et al. 2010; Heckmann et al. 2012), where consumers change the targets or strengths of their trophic interactions to acquire better foraging payoffs in response to changing resource abundances or other conditions. Optimal foraging (OF) theory provides a potentially powerful mechanistic basis for adaptive foraging that considers energetic payoffs of consumers when foraging under different resource availabilities (Charnov & Orians 1973). A consumer following OF feeds on a specific subset of its potential resources that maximises its energy intake given the current resource abundances. This subset (the optimal diet) is context-dependent and subject to change as resource abundances vary.

Consistent with the potential importance of diet choice, past work has shown that OF is able to partly predict realistic food-web structures (Beckerman et al. 2006; Petchey et al. 2008; Thierry et al. 2011b; Ho et al. 2019). However, these studies have mostly applied OF as a static, one-time determination of species’ diets. In contrast, the context-dependent manner in which OF is formulated (Charnov & Orians 1973; Krebs et al. 1977) means that consumers should ideally adjust their optimal diet choice dynamically in response to changing resource abundances. As a result, food-web structures generated through OF should also not be fixed, but instead dynamically changing (Beckerman *et al*. 2010; Loeuille 2010), as has been observed empirically (McCann & Rooney 2009; Ushio et al. 2018).

Beyond changes in food-web structure, the impacts of OF on population dynamics in food webs may influence whether species can persist over time, which has important conservation implications (Tylianakis et al. 2010). Intuitively, we may presume that adaptive foraging is a stabilising mechanism, since by definition it benefits the consumers and should thus help them to persist (Rooney *et al*. 2006; Heckmann et al. 2012). Yet, this is not necessarily the case when viewing the food web as a whole, because any species’ advantage could be disadvantageous to others in an antagonistic interaction system. In this respect, the effects of adaptive foraging on species persistence in complex food webs have mostly been theoretically studied through interaction-strength adjustments (e.g., reallocation of foraging effort, Kondoh 2003, 2006; Uchida et al. 2007; Guill & Drossel 2008), or an instant rewiring that happens only following species extinctions (e.g., Thierry et al. 2011a). However, the effects of a continuous diet rewiring mechanistically driven by OF has not yet been considered. More importantly, while classical OF theory assumes that consumers perceive and react to changing resource abundances perfectly (Charnov & Orians 1973; Krebs et al. 1977), in reality, there are inherent (e.g., sensory or cognitive) limitations that prevent such perfect responses. Indeed, past work based on the simplest one-consumer-two-resource system has shown that the consumer’s optimal diet choice with different effectiveness (i.e., levels of optimisation) can lead to distinctive dynamical outcomes Fryxell & Lundberg (1994); Mougi & Nishimura (2009), indicating that the effectiveness of diet adjustment is likely an important factor influencing the food-web consequences driven by OF. Yet, studies on this effectiveness have been limited to the above-mentioned simple systems, whereas its influences in complex food webs remains unknown.

Here we aim to fill these knowledge gaps by simultaneously investigating how community species persistence and food-web structures respond to the mutually-dependent OF diet adjustment and population dynamics. In doing so, we consider deviations from perfect OF diet adjustments by comparing schemes with the same initial state but different levels of diet-adjusting effectiveness. We ask whether OF-based adaptive foraging can stabilise food webs by (i) making them more resilient, i.e., dampening structural variation or allowing structures to re-establish after species loss, and (ii) improving overall species’ persistence. Furthermore, we pay particular attention to how effectiveness of diet adjustment influences these OF-driven food-web consequences.

## Methods

### Overview

We simulated a pool of species with empirically realistic body masses and abundances, and then drew a 50-species synthetic community (*×*30 replicates) from the pool. We fixed the smallest five species (in body mass) to be basal resources while the remaining consumers, to mirror empirically observed basal-resource proportions and large-eat-small tendency (Ho et al. 2021). Each consumer species could potentially consume all other species except itself (i.e., cannibalistic links were prohibited). We then “wired” these species into a synthetic food web based on diet predictions derived at their given abundances using an optimal foraging model. Taking this food web and the corresponding species abundances as the initial state, we simulated species population dynamics using a generalised Lotka-Volterra model under schemes that differed in the effectiveness (i.e. rate) of optimal foraging diet adjustment, quantifying food-web structural measures and species extinctions during the course. We thus compared food-web consequences among foraging schemes emerging from a shared initial state. All simulations and analyses were done in R version 3.4.4 (Team et al. 2013). Details about the models and simulations are given in the following subsections.

### The optimal foraging model

Optimal foraging (OF) depicts consumers as always selecting the optimal diet, which is constituted of a specific subset of the pool of potential resources that maximises the consumer’s energy intake rate. This intake rate necessarily depends on the energy contents of resource items, the consumer’s encounter rates with resource items (accounting for resource abundances), and the consumer’s handling time on resource items, modelled by a multi-species Type-II functional response (Holling 1959; Drossel & McKane 2005; Ho et al. 2019). Here, we assume that a unit biomass of any resource species provides the same amount of energy to any consumer species. The consumer’s energy intake rate can thus be evaluated by its total (i.e., sum over its diet) biomass consumption rate.

In a food web, where multiple species are feeding on each other, the total biomass consumption rate of the *j*^th^ consumer species is

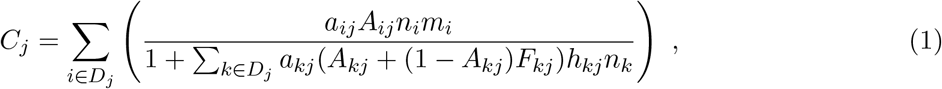

where *D*_*j*_ is the set of *k* species that constitutes its diet, and for the *i*^th^ resource species in its diet, *a*_*ij*_ is its per-capita search rate (m^2^s^*−*1^ as we model 2D foraging), *A*_*ij*_ its attack success probability, *h*_*ij*_ its per-capita handling time (s) of a successful foraging attack, and *F*_*ij*_ quantifies its time cost for engaging in an unsuccessful attack attempt (*sensu* Meire & Ervynck 1986), expressed as a proportion (here assumed to be a constant) of *h*_*ij*_. *n*_*i*_ is the *i*^th^ resource species’ numerical abundance (here, 2D density, in m^*−*2^) and *m*_*i*_ its body mass (kg; mean body mass ignoring intraspecific variation). For detailed parameterisation, see Supplementary Information section S1.

To identify a consumer’s optimal diet, one needs to find the specific set *D* that maximises its resultant *C* with given resource abundances. In response to changing resource abundances, the diet adjustment predicted by OF—either including new resources into, or excluding existing resources from, the current OF diet—must always follow the profitability order of the consumer’s potential resources (Ho et al. 2019). Profitability is the expected energy (here, biomass) gain per time of an already-encountered resource item, which is independent from the resource’s abundance. Dividing the expected biomass gain by the expected time spent on attacking, the *i*^th^ resource species’ profitability to the *j*^th^ consumer species is therefore

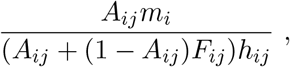

Once resource profitabilities have been calculated, the consumer’s realised optimal diet can then be derived by identifying the highest *C* generated with eqn (1) from its narrowest (i.e., eating only the most profitable species) to broadest (i.e., eating all species) possible diets following a decreasing profitability order. The above OF model essentially builds on the “Allometric Diet Breadth Model” (Petchey et al. 2008) by developing further its mathematical structure and applying updated mass-scaling constraints on parameters.

### Population dynamics

The population dynamics of species in each focal food web were modelled using a generalised Lotka-Volterra model, where the species’ numerical abundances described by ordinary differential equations as (in matrix notation)

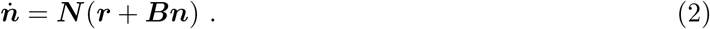

Here, ***r*** and ***n*** are vectors of species’ intrinsic growth rates (*r*) and numerical abundances (*n*), respectively. ***B*** is the *S × S* (*S* = species richness) interaction matrix with trophic interaction strengths as off-diagonal entries (0 if no interaction) and intraspecific interference strengths as diagonal entries. ***N*** is a *S × S* matrix with the values of *n* along its diagonal and 0 elsewhere.

Following previous work (Pawar et al. 2012; Ho et al. 2021; Tang et al. 2014), we assume that a species’ intrinsic growth rate and intraspecific interference strength both scale with its body mass. The per-capita intrinsic growth rate of the *i*^th^ species scales as

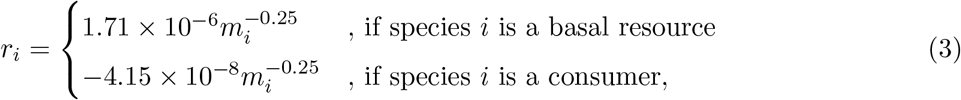

where *m*_*i*_ is its body mass (kg). The per-capita intraspecific interference strength of the *i*^th^ species (*b*_*ii*_, the diagonal elements of ***B***) scales as

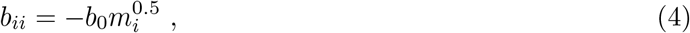

where *b*_0_ a positive scaling constant (for values, see further below). The negative intrinsic growth rates represent consumers’ natural mortality rates, while the negative interference strengths represent the typically negative effects of increasing density on species’ population growth.

The per-capita trophic interaction strengths between the *j*^th^ consumer species and its *i*^th^ resource species (coupled *b*_*ij*_ and *b*_*ji*_, the off-diagonal of ***B***) can then be derived from biomass consumption rates by accounting for the fraction of population numerical loss and gain contributed by per-capita consumer and resource, respectively. That is,

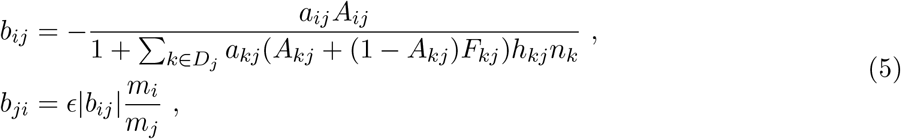

The first equation is the numerical loss from the population due to being consumed, and the second the numerical increase from consuming others. The scalar *1* is the consumer’s efficiency of converting consumed biomass to its own biomass (here assumed to be a constant, see SI section S1).

### Simulation of food-web dynamics

Building a synthetic species pool, we simulated 100,000 species whose body masses (kg) were randomly drawn from a log-normal distribution (Preston 1948; Engen & Lande 1996; Ho et al. 2019) that covered the range 10^*−*7^–10^3^. Their abundances (densities) were assigned based on a body-mass scaling rule (“Damuth’s law” Damuth 1981; Brown et al. 2004) such that the numerical abundance (*n*) of the *i*^th^ species is

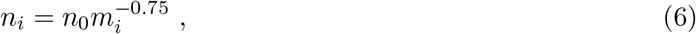

where *m*_*i*_ is its body mass, and *n*_0_ a positive scaling constant (for value, see further below). Synthetic 50-species communities for establishing food webs were then drawn from the pool for each of the 30 replicates. Because none of our hypotheses pertained to priority effects or invasion, we set the communities with all 50 species simultaneously, though we acknowledge that sequential additions of species could give different outcomes (e.g., Drossel et al. 2001).

We wired all synthetic communities into “initial webs” for subsequent simulation of population dynamics. An initial web could be set at a steady state where all species coexist with positive abundances, and all consumers’ current diets are exactly their optimal diets derived (eqn (1)) with such species abundances. However, an steady state cannot be analytically solved *a priori* in a Type-II system with its dynamic abundance-dependent interaction strengths, and a fully feasible (i.e., all species coexist) equilibrium is highly improbable with all metabolic constraints (i.e., mass-scaling rules) we have set. Therefore, we approximated such a steady state, and ensured that all generated initial webs are comparable, by constraining each initial web to have the following properties:

- The species abundances are as close as possible to an “analytical” dynamical equilibrium of the system (solved assuming fixed interaction strengths), yet still follow the body-mass scaling rule (eqn (6)) and thus all abundances must be positive. The chosen analytical equilibrium, among all possibilities, should therefore be the most feasible one, i.e., having as many species exhibiting positive equilibrium abundance as possible (see Ho et al. 2021).
- All consumers feed on their optimal diets based on the given values of species abundances and all other parameters, i.e., the initial web is itself an OF web. This ensures that, across replicates, any OF diet adjustment taking place during simulation is triggered by species abundance fluctuation after the simulation begins, not by any preset difference among initial webs.
- The web has a ∼ 0.1 connectance, which is comparable to observed values in empirical food webs (Dunne et al. 2002a; Ho et al. 2019). All initial webs across replicates thus have not only a comparable number of nodes (i.e., 50 species) but also number of links.

These criteria were imposed by finding the suitable set of scaling constants of species abundance (*n*_0_, eqn (6)), handling time (*h*_0_, see SI section S1), and interference strength (*b*_0_, eqn (4)), while still complying with the size-scaling rules (by keeping the scaling exponents fixed). The connectance of an OF web can be controlled by adjusting values of *n*_0_ and *h*_0_ for all species, given that a consumer’s optimal diet breadth becomes narrower with increasing abundances of profitable resources or increased handling times (Ho et al. 2019). For each synthetic community, we generated OF webs using eqn (1), by first setting the *n*_0_ of all species sequentially between 10^*−*6^ to 10^2^ (with a 0.2 increase in the exponent each time), and then calculated the corresponding *h*_0_ that guaranteed the target connectance value of 0.1 of the web. For all these generated OF webs, we then assigned species’ interference strengths by varying the values of *b*_0_ between 10^*−*8^ and 10^8^ (with a 0.2 increase in the exponent each time). With the assigned interference strengths (eqn (4)), as well as the known trophic interaction strengths (eqn (5) with above-given *n*_0_ and *h*_0_ combinations) and intrinsic growth rates (eqn (3)), these OF webs’ equilibrium abundances (***n***^**\***^) can be calculated by solving eqn (2) as

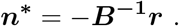

Notably, these are just analytical solutions for any given set of fixed parameter values, which could include negative equilibrium abundances (biologically insensible) and do not reflect true outcomes of OF-driven population dynamics. They were here calculated for us to pick an appropriate reference to decide an initial web. To achieve this, we then focused on the *b*_0_ that gives the most feasible equilibrium to each of the *n*_0_ and *h*_0_ combinations. Among these most-feasible equilibria across *n*_0_ and *h*_0_ combinations, the one with species abundances scale closest to Damuth’s law with a −0.75 mass-scaling exponent, judging by the smallest residual of a linear regression on species body masses to their abundances (both in log_10_ scale), was identified and taken as the reference equilibrium. The specific parameter set of *n*_0_, *h*_0_, and *b*_0_ leading to this reference equilibrium, the mass-scaled species abundances with this *n*_0_ (not the reference equilibrium abundances), and the corresponding OF web wired at their combination (i.e., the initial web) together presented the “initial state” for subsequent food-web dynamical simulation.

We then simulated the population dynamics from the initial state by numerically solving eqn (2) using the `deSolve` R package (Soetaert et al. 2010) and its `ode` function. Each simulation was run for 10^8^ time units (10^3^ as an integration time step). During the simulation, any species whose abundance (density) fell below 10^*−*9^ was considered extinct and its abundance set to 0. From the same shared initial state, food-web dynamics were allowed to unfold based on each of five foraging schemes (so five separate simulations per initial state):

- Type.I: the trophic interaction strengths are fixed (i.e., not resource abundance-dependent) to values matching the initial state, and the consumers do not adjust their diets at all over the simulation. In other words, this scheme essentially simulates dynamics with Type-I functional responses (although the initial-state interaction strengths are themselves derived with a Type-II function as eqn (1)). Note that as species abundances at the initial state are close, yet never equal, to those at the reference analytical equilibrium, population dynamics with abundance fluctuations will still occur in this scheme even with fixed interaction strengths. This is a “null”, control scheme that eliminates both the abundance dependence of parameters and the consumers’ OF diet adjustment.
- Type.II: the trophic interaction strengths are dynamically updated with real-time species abundances following eqn (5), but the consumers are still not adjusting their diets. This can be seen as another null scheme, simulating dynamics with Type-II functional responses but without consumers’ OF diet adjustment.
- OF.fast: the trophic interaction strengths are as in the Type.II scheme. The optimal diet of each consumer is re-calculated every 10^4^ time units (10 time steps) based on species abundances at that juncture using eqn (1). Consumers are accordingly allowed to adjust their diet by one resource, following the resources’ profitability order towards achieving their currently optimal diet. That is, a consumer may broaden or narrow its diet by one resource, or keep the same diet, depending on which act makes its diet closer to the re-calculated optimal one. The population dynamics simulation is resumed after the diet adjustment with an updated food-web topology and corresponding interaction matrix (***B***).
- OF.mid: the same as the OF.fast scheme, but the re-derivation of optimal diets and consumers’ diet adjustment happens every 10^5^ time units (100 time steps).
- OF.slow: the same as the OF.fast scheme, but the re-derivation of optimal diets and consumers’ diet adjustment happens every 10^6^ time units (1000 time steps).

We expect the comparison of the Type.I and Type.II schemes to reveal the influence of resource abundance-dependence of trophic interaction strengths. Comparing the Type.II with the OF schemes is expected to reveal the influence of OF diet adjustment. Specifically, by not allowing an instantaneous and perfect diet adjustment of consumers to their respective optimal diets, but instead imposing an incomplete switch (at three frequency levels: OF.fast, OF.mid, and OF.slow), we aim to quantify the effect of different diet-adjusting effectiveness. The more frequently consumers are allowed to adjust in a certain period of time, the more effectively they can keep up with the fluctuating resource abundances and better approach their optimal diets.

We recorded species extinctions (at every 10^3^ time units) and food-web structural properties (at every 2.5 *×* 10^7^ time units) during the dynamical simulation. The following structural properties were recorded: connectance, number of top consumers, nestedness, modularity, and mean (realised) resource-consumer body-mass ratio (in log_10_ scale). Their definitions and measures are provided in SI section S2. We compared cumulative species extinctions and patterns of fluctuations of these structural features through time among the five schemes.

## Results

In terms of both species persistence and food-web structure, our simulation generated clear scheme-specific patterns. As species abundances in the initial state were chosen to be close to an analytical equilibrium of the (initial) web for a given set of parameter values, in the Type.I scheme, where trophic interaction strengths were fixed, there were indeed rather few species extinctions (Fig. 1) over time. In contrast, in the Type.II scheme, where trophic interaction strengths changed with fluctuating abundances, extinctions happened relatively frequently (Fig. 1). As neither of these schemes allowed consumers to carry out OF diet adjustment, and their dynamics started from exactly the same state, the more extinctions in the Type.II than in the Type.I scheme were caused purely by the abundance-dependent variation in trophic interaction strengths driven by the consumers’ Type-II functional responses in the former.

**Figure 1:**
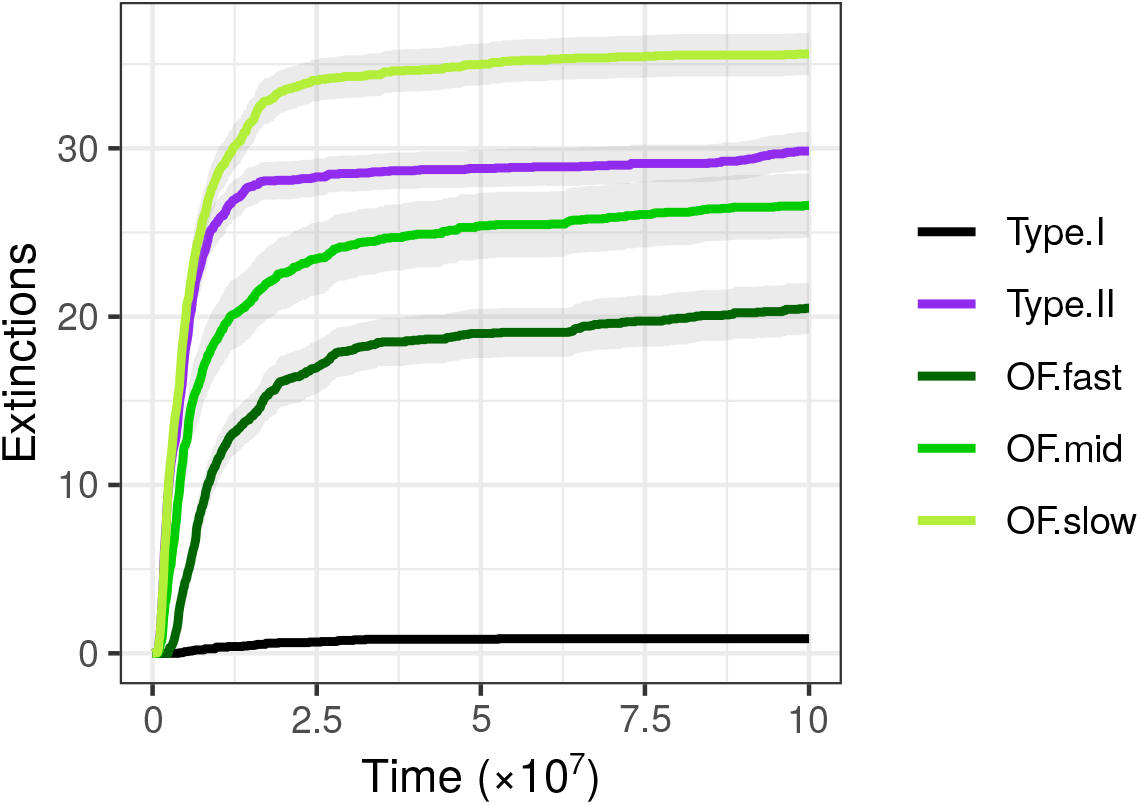
Cumulative extinctions over time in food webs (N = 30). The solid lines indicate the mean while the shaded areas present the standard error. Taking the Type.II as the reference, the OF.fast and OF.mid schemes, with their relatively effective OF diet adjustment, saw fewer cumulative extinctions. In contrast, the OF.slow scheme, with its relatively ineffective diet adjustment, saw more cumulative extinctions.

The OF.fast and OF.mid schemes had fewer cumulative extinctions than the Type.II scheme, indicating that adaptive diet adjustment mitigated some of the species extinctions caused by Type-II functional responses. However, this mitigating effect was reversed when consumers’ diet adjustment was not effective enough, as the OF.slow scheme had even more cumulative extinctions than the non-diet-adjusting Type.II scheme (Fig. 1). Notably, as we judged species extinction using an abundance threshold (*Methods*), in the Type.II scheme there were occasionally a few consumers having all their initial resources go extinct without yet having their own abundances drop below the threshold by the end of the simulation. These “resourceless” consumers were actually doomed to go extinct without the ability to establish new links to resources, but were technically not recorded as extinctions— the number of extinctions in the Type.II scheme was thus slightly underestimated. Therefore, more precisely speaking, the OF.slow scheme may not necessarily have led to more extinctions (Fig. S1), but certainly accumulated extinctions faster (i.e., higher extinction rates) than the Type.II scheme. Ineffective OF diet adjustment, though by definition being still adaptive (in terms of gaining more energy) to individual consumers, was surprisingly harmful to the overall species persistence in food webs.

Food-web structure also varied through time. As no diet adjustment was allowed in the Type.I and Type.II schemes, all their structural changes were purely driven by the loss of nodes and links via species extinctions during the dynamics. Accordingly, the Type.I scheme saw very little structural change throughout, while the structure in the Type.II scheme first changed rapidly then became near-constant (Fig. 2), largely reflecting the patterns of cumulative species extinction (Fig. 1). In general, food webs in the Type.II scheme became less connected, less nested, and less modular over time. Also, their number of top consumers slightly decreased, while the mean resource-consumer body-size ratio slightly increased (Fig. 2). Conversely, food webs in the OF schemes showed different structural change trajectories. Generally, these food webs became slightly more connected, equally nested, less modular, had fewer top consumers (even fewer than the in the Type.II schemes) and a slightly smaller mean resource-consumer body-size ratio by the end of dynamical simulation (Fig. 2). Notably, unlike in the Type.I and Type.II schemes, here in the OF schemes the structural changes were driven by both species extinction and trophic link rewiring due to consumers’ diet adjustment. Therefore, food-web structures did not change toward their final states monotonically, but instead had fluctuations throughout the simulation (Fig. 2). Looking at the ending phase of simulation, in most structural measures, the OF schemes tended to be closer to the shared initial state than the Type.II scheme, and the OF.fast tended to be the closest (Fig. 2). These findings suggested that, while adaptive diet adjustment is itself a mechanism that can alter food-web structure, an effective diet adjustment can, in the long run, maintain some structural properties even though the size of the food web may have changed due to species extinctions.

**Figure 2:**
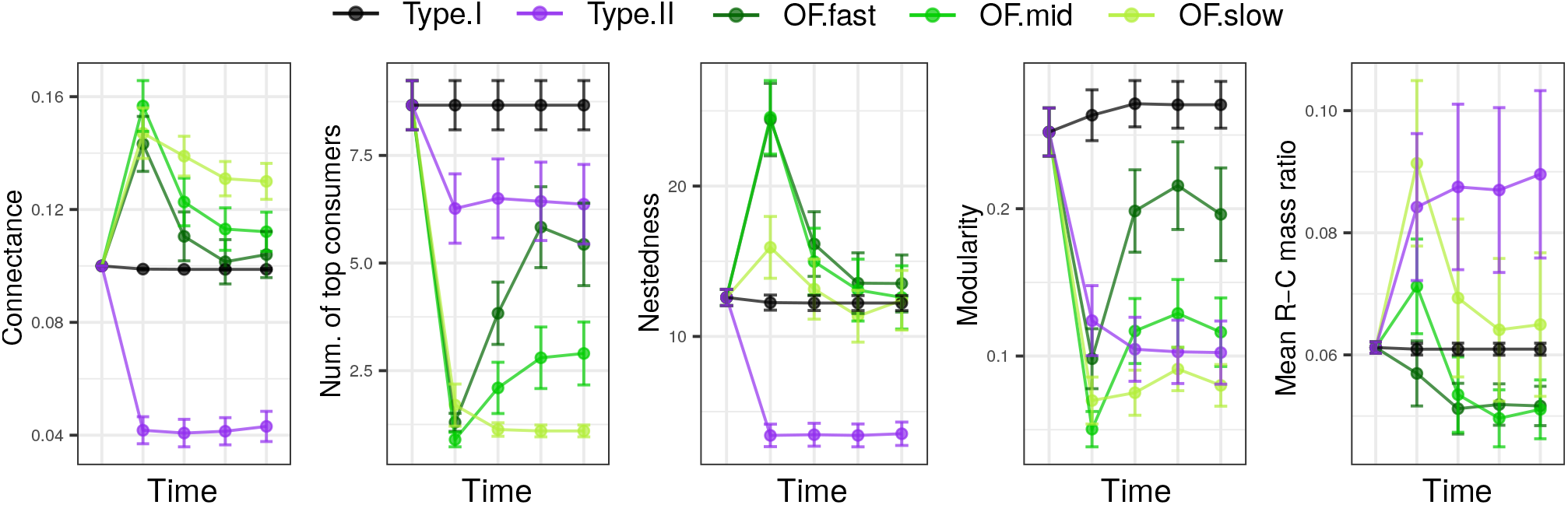
Structural measures of the food webs over time (N = 30). Points indicate the mean while the error bars present the standard error, given at the five time points tick-marked in Fig. 1.

A closer look at species identities within food webs (Fig. 3) further revealed the qualitative difference in extinctions (i.e. “who went extinct”) among the schemes. This also helped us to better understand the causal links between diet change, extinctions, and changes in food-web structure. In the Type.II scheme, the initial extinctions could occur on consumer species of any body size (Fig. 3), but most large consumers went extinct eventually, leaving some intermediate- and small-sized consumers in the food web by the end of simulation. Although species were not adjusting their diet in the Type.II scheme, due to the extinction of large consumers, some intermediate- or small-sized consumers were freed from predation and themselves became new top consumers (Fig. 3). As a result of such trophic role transitions, the decrease in number of top consumers did not appear to be severe (Fig. 2) even though the “initial” top consumers had mostly gone extinct. In contrast, the OF schemes immediately generated species role transitions through rewiring rather than extinction (Fig. 3). Early in the dynamics, the large consumers broadened their diet toward smaller species (SI Fig. S2), making the web more connected and more nested while less modular. Meanwhile, the smallest few top consumers became the resources of larger ones, thus their roles turned into intermediate consumers and caused a decrease in the number of top consumers (Fig. 2, 3). Over time, species extinction was largely suppressed in the OF.fast schemes, where those that went extinct were mostly large-sized intermediate consumers. As diet adjustment became increasingly ineffective in the OF.mid then the OF.slow scheme, increasing numbers of smaller-sized intermediate consumers went extinct (Fig. 3).

**Figure 3:**
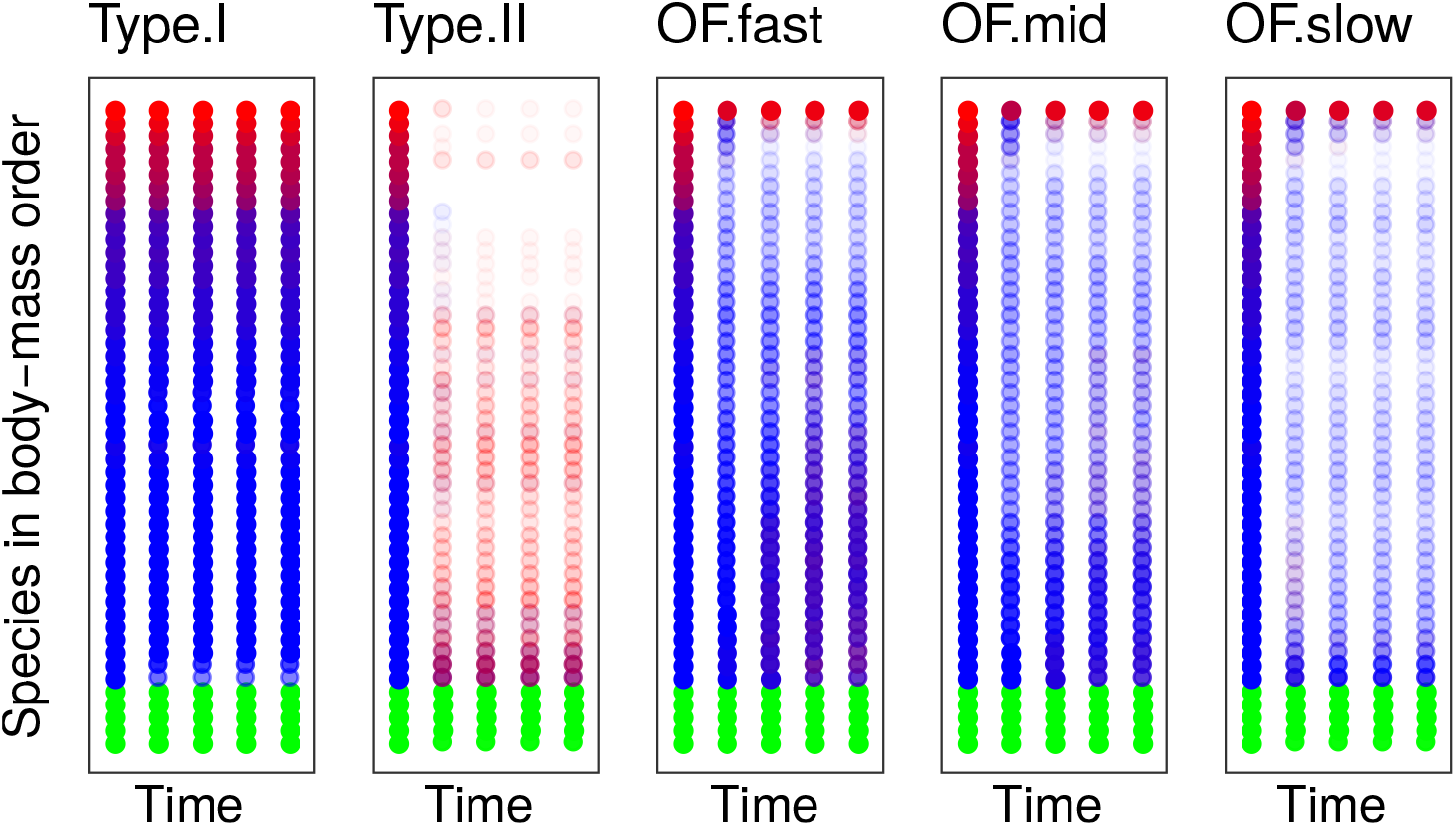
Species body masses, trophic roles, and dynamical fate (go extinct or not) within food webs (N = 30). The five columns within each scheme panel, from left to right, correspond to the five time points as tick-marked in Fig. 1. The red/blue/green colours indicate the trophic roles, i.e., top consumer/intermediate consumer/basal resource, respectively. The white colour indicates that the species went extinct (including isolated “resourceless” consumers, when present, in Type.I and Type.II). As each nodes’ status is averaged over 30 replicates, a stronger colour fading (becoming white) indicates that a greater proportion of species at this body-mass rank went extinct.

## Discussion

Understanding the mechanisms that shape the structural and dynamical properties of ecological networks is necessary for resolving the complexity of nature, and is the foundation for dealing with critical ecological issues regarding biodiversity and ecosystem functioning (Dunne et al. 2002b; Tylianakis *et al*. 2010). We have shown that organism-level adaptive diet choice behaviour, based on optimal foraging (OF), indeed has strong structural and dynamical consequences for food webs. Particularly, how effectively species perform such diet adjustments is a crucial factor determining whether adaptive foraging benefits or harms the overall species persistence in food webs.

Our simulations showed that abundance-dependent trophic interaction strengths following Type-II functional responses, in comparison to fixed strengths, exacerbated species extinctions (Fig. 1). This is expected, as the Type-II functional response has long been known to negatively affect species persistence in consumer-resource systems (Oaten & Murdoch 1975). Mathematically, under the governance of Type-II functional responses, the per-capita consumption pressure that a consumer imposes on a resource species is positively associated with the resource’s abundance (Dunn & Hovel 2020; Fig. 4). Therefore, consumption suppresses the resource’s population growth when it is rare, but facilitates its growth when abundant—this kind of feedback amplifies species population fluctuations and is exactly the opposite of what is required to allow species persistence. In contrast, OF diet adjustment can (to a degree) counteract the negative effect of Type-II interaction strengths on species persistence because under this foraging regime, when the currently-consumed resource becomes rare, the consumer tends to broaden its diet to include other less-profitable resources, such that the per-capita predation pressure on the currently-consumed resource will be diluted (Rooney et al. 2006). This dilution provides the feedback required for species persistence (Fig. 4), and we therefore observed fewer cumulative extinctions in the OF.fast and OF.mid schemes than in the Type.II scheme (Fig. 1).

**Figure 4:**
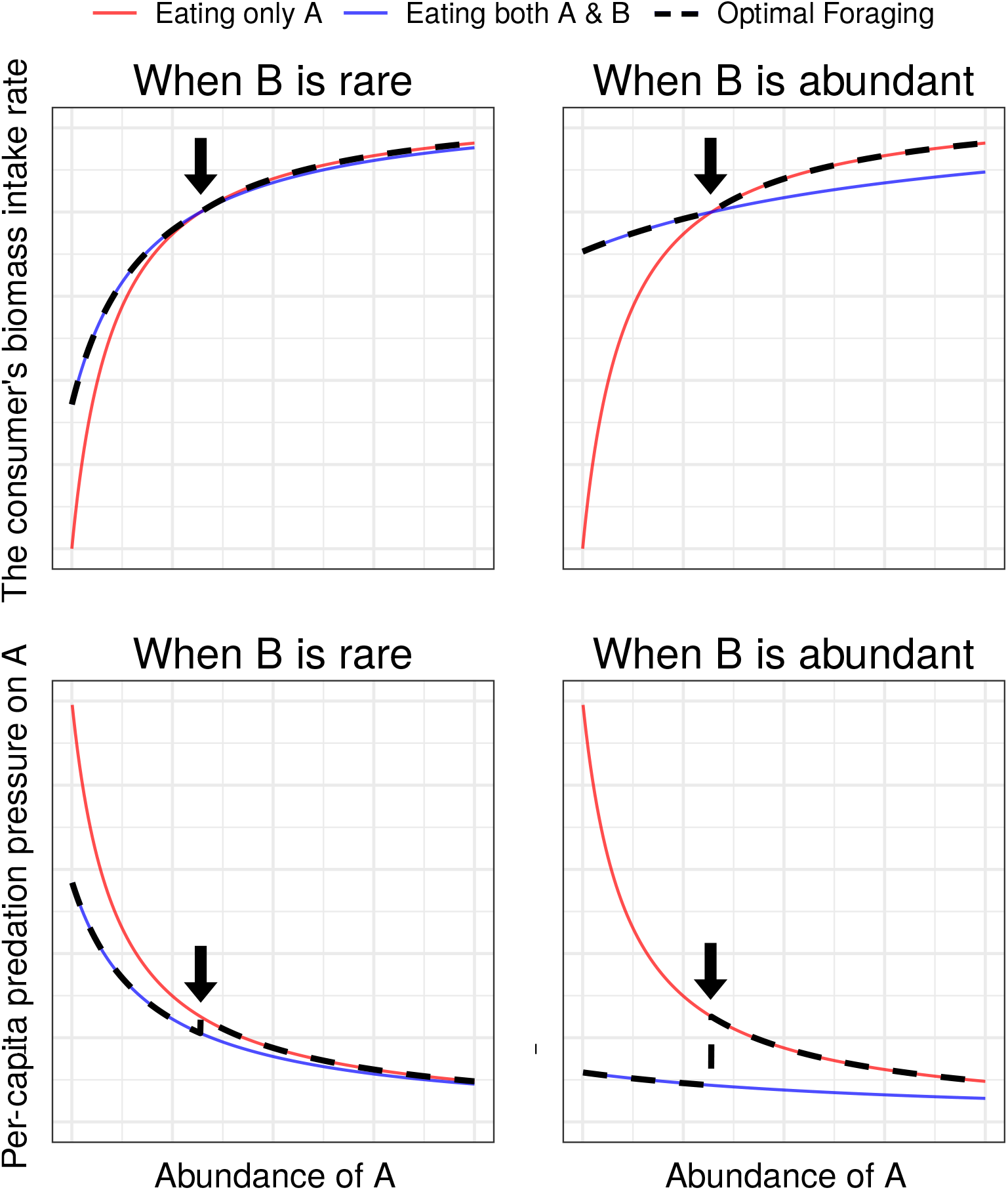
A consumer’s biomass intake rate and the predation pressure it imposes on percapita more-profitable resource. This is illustrated in a one-consumer-two-resource system. A and B are the two resource species, where A is the more-profitable one. The black dashed lines indicate the scenario with optimal foraging diet adjustment. The biomass consumption rate (top panels) follows a multi-species Type-II function, and the per-capita predation pressure on A (bottom panels) is quantified by A’s contribution to the consumer’s biomass intake rate divided by A’s numerical abundance. The consumer that forages without diet adjustment but feeds “single-mindedly” on A (red lines) leads to a monotonically increasing predation pressure on A with its decreasing abundance (bottom panels), which gives the typical destabilising effect of a Type-II functional response on A’s population. In contrast, making a diet adjustment to include B when A is rare (black dashed lines) creates a sudden fall of predation pressure at the threshold abundance of A (indicated by arrows, bottom panels). This leads to a lower predation pressure at the low-abundance end of A than in the previous scenario, and therefore mitigates partly the destabilising effect. Comparing the left with right panels reveals that such mitigation is more effective when the less-profitable resource (B) is abundant.

Interestingly, we found that OF diet adjustment increases species persistence only when the adjustment is frequent, and thus effective, enough; otherwise, diet adjustment surprisingly intensifies species extinctions (see comparison of OF.slow vs. Type.II scheme, Fig. 1). Rooney *et al*. (2006) have previously shown that rapid prey switching of predators at higher trophic levels is needed for regulating the consumption pressure among resources and stabilising different resource channels in the system. Therefore, the too-ineffective diet adjustment of consumers—by failing to keep up with the pace of resources’ population dynamics—may lead to a mismatch between the consumption pressure and the abundance of their resources. The resultant disproportionately high pressure on species at lower trophic levels, especially on those remaining after some extinctions have occurred, may facilitate subsequent extinctions of resource species. Indeed, most extinctions in the OF.slow scheme were of intermediate but not top consumers (Fig. 3). While most previous studies have suggested that adaptive foraging positively contributes to species persistence in food webs (e.g., Kondoh 2003; Heckmann et al. 2012), some reported the opposite (Gilljam et al. 2015; Gauzens et al. 2021). Our findings help to reconcile these conflicting results: both outcomes are possible depending on the effectiveness of the adaptive-foraging adjustment. These previous studies were sufficiently distinctive in their approach to modelling adaptive foraging and population dynamics that their results may not be directly comparable. Nevertheless, a manipulation of adaptive-foraging “effectiveness” relative to the pace of population dynamics can certainly be incorporated into various models. Furthermore, the key role of adaptive-foraging effectiveness can likely be tested even with relatively simple experiments. For instance, with well-monitored food webs in mesocosms where the consumers’ cognition of resource abundances, and thus the information basis of performing effective adaptive foraging, is manipulated. Based on our findings here, we would hypothesise that higher extinction rates can be detected when the consumers’ cognition is sufficiently suppressed or misinformed. Checking whether effectiveness is the determinant of positive vs. negative effect of adaptive foraging on species persistence, in both yet various forms of theoretical and empirical food webs, would be an fruitful future direction.

By mapping species’ body masses and trophic roles to the extinction outcomes, we find that not all species have the same extinction risk. Basal resources, with their positive intrinsic growth rates set in the population dynamics model, mostly persist. In the long run, large consumers are more susceptible to extinction (Fig. 3). Although the OF diet adjustment allows the very few largest consumers to persist. The remaining large consumers besides these few still eventually go extinct, even with relatively effective diet adjustment (Fig. 3). Notably, again echoing Rooney *et al*. (2006), the ability of large consumers to quickly adjust their diets seems to be particularly important for the system to avoid early extinctions caused by Type-II functional responses (OF.fast & OF.mid schemes vs. Type.II & OF.slow schemes in SI Fig. S2). Therefore, body size seems to play a key role in influencing the contribution of OF diet adjustment to species persistence in size-structured food webs. From an ecological perspective, species with different body sizes exhibit distinctive life-history traits (Brown et al. 2004). Large species have relatively low population growth potential, so their numerical population response to changing resource abundances would be ineffective, and large individuals in their relatively long life span would likely experience multiple fluctuations of their resource abundances. Thus, it is reasonable to hypothesise that large species should have evolved effective behavioural diet-adjusting abilities to better respond to resource fluctuations. Indeed, making behavioural diet adjustment requires good cognitive abilities for assessing the changing environment and finding resource patches, as well as a suitable set of traits (e.g., strong-enough body and multi-functional mouth part) to be able to feed on alternative types of resource. These are also features more likely being possessed by larger species (Mittelbach 1981; Costa et al. 2008). Putting together our simulation results and these ecological considerations, and given that body size is a major determinant of animal behaviour (e.g., Cozzoli et al. 2019), we hypothesise that there likely exists an “allometry” of diet-adjusting effectiveness in the real world (Dial et al. 2008), presumably scaling positively with species body mass, i.e., large species are better able to achieve effective diet adjustment. Extending our current theoretical framework, perhaps with explicit species-specific generation-time settings, to further investigate this potential diet-adjusting allometry is a promising avenue for a next-step research.

Food-web structures have traditionally been considered to have been shaped in two ways: by lower-level mechanisms (e.g., food-web nestedness emerges from body-size constraints of foraging, Coelho et al. 2021) or by whole-system stability (e.g., modularity confines cascading extinctions so modular food webs can persist, Stouffer & Bascompte 2011). Both factors are valid, with the realised structures of food webs reflecting their combined constraints (*sensu* Ho et al. 2021). With this perspective, we defined distinct behavioural foraging regimes, letting the structure and population dynamics of the food web emerge in each regime. In other words, we allowed both the lower-level and whole-system factors to operate during the dynamics. The species persistence and food-web structural patterns we have captured are therefore the dynamical consequences shaped by the two factors combined. Such a consideration of both factors at the same time can be a key step toward a better comprehension of how structural signatures of real-world food webs are formed.

Our understanding of empirical food webs has been, so far, largely based on temporal snapshots (but see Schmidt et al. 2009). Each snapshot reflects one—but not necessarily the only—state of the community. In nature, abundances of species typically fluctuate (e.g., Wolda 1978; Colebrook 1985), thereby influencing the rewards associated with the use of any given species as a resource. Abundance-driven adaptive foraging thus makes food-web structure a property that changes over time, and feeds back to species’ population dynamics. Indeed, the trajectories of structural changes revealed in different schemes of our simulations (Fig. 2) imply that food-web structure fluctuates, and the degree to which it varies depends strongly on the underpinning foraging behaviours. Inspired by these findings, an interesting follow-up question would be: do we observe the same association between adaptive-foraging effectiveness and structural variations in real-world food webs? Besides the manipulative experiments as above-mentioned, this question can potentially be examined by comparing temporal food-web structure among systems where adaptive foraging may intrinsically be constrained to different degrees. To elucidate such an idea: in this study, for the sake of simplicity and generality, we did not specify any realistic biological identity of our synthetic species, but assumed that any consumer can potentially feed on any other species in the community, and body size is the one trait that determines foraging parameters and eventually the realisation of trophic links. Nonetheless, we recognise that other traits (or say, trait-matching between consumers and resources) are also prevalent determinants of trophic interactions in nature (Eklöf et al. 2013). In cases where trophic interactions are strongly constrained by certain traits other than body size, these traits as edibility filters would prohibit a consumer from feeding on some resources. The consumer would thus have fewer potential resources, thus a shrunk capability to make diet adjustments and a suppressed adaptive-foraging effectiveness. Extrapolating the idea, we would hypothesise that adaptive foraging is generally more effective in systems where trophic interactions are mainly driven by “general” traits (e.g., body size in aquatic food webs; Potapov et al. 2019), and there we expect to see smaller food-web structural fluctuations, whereas the opposite in systems driven by “specific” trait-matching (e.g., chemical tolerance in herbivore-plant food webs; Agrawal et al. 2012). Indeed, studying adaptive foraging in real-world complex food webs is challenging and thus rather rare, as one would need time series highly-resolved data to observe how rewiring actually happens. To our knowledge, nonetheless, there are some relatively long-term, repeatedly sampled empirical food-web data recently becoming available (Schmidt et al. 2009; Ushio *et al*. 2018). Further accumulation of such data would favour the implementation of the comparisons suggested above. As a food web’s structure determines the functions that it provides (e.g., Setälä et al. 2005), knowing if and how food-web structure changes over time is essential for estimating the temporal variation of ecosystem functions, and is arguably a more thorough approach than those purely informed by species richness or abundances.

In conclusion, the ability to make adaptive diet adjustments in response to fluctuating resource abundances benefits the consumers and stabilises the whole food web in terms of its structure and overall species persistence, but only when consumers’ diet adjustment is sufficiently effective. Ineffective diet adjustment, instead, aggravates extinctions in the food web, especially of intermediate consumers. The effectiveness of making behavioural adjustment is therefore a determinant of the effects of adaptive foraging on communities but has not been examined mechanistically until now. Our findings thus reveal a novel aspect of organism-level foraging behaviour that can scale up to shape community-level properties. Focusing on the energetic basis of organismal behaviour as the fundamental driver of ecological network formation and dynamics can enlighten paths toward a better mechanistic understanding of complex ecosystems.

## Acknowledgements

We thank Andrew Beckerman, Robert Ewers, Axel Rossberg and Erol Akcay for their comments on earlier drafts of this manuscript. HH was funded by Imperial College President’s PhD Scholarship. SP was supported by Natural Environment Research Council (NERC) grants NE/M020843/1 and NE/S000348/1. JMT was funded by the Marsden Fund (grant number UOC1705).

