## Supplementary information for "Less is worse than none: ineffective adaptive foraging can destabilise food webs"

Hsi-Cheng Ho[1, a], Samraat Pawar[1], and Jason M. Tylianakis[2]

[1]Department of Life Sciences, Imperial College London, Silwood Park Campus, Ascot SL5 7PY, UK
[2]Bioprotection Aotearoa, School of Biological Sciences, University of Canterbury, Private Bag 4800, Christchurch, NZ
[a]Current Address: Department of Aquatic Ecology, Eawag: Swiss Federal Institute of Aquatic Science and Technology, Dübendorf 8600, CH

### Contents

**S1 Parameterisation** **2**

**S2 Food-web structural measures** **4**

**S3 Additional figures** **5**

**S4 Additional discussion** **7**

### S1 Parameterisation

Here we describe how foraging parameters of the optimal foraging diet-choice model (eqn (1) in the main text) and the conversion efficiency in the population dynamics model (eqn (5) in the main text) are specified. All quantities are in per-capita SI units.

For the conversion (or assimilation) efficiency of consumers ($\epsilon$), we assume it is approximately constant, irrespective of the consumer and resource identities (Kondoh 2007; Petchey *et al.* 2008) or their body sizes (DeLong *et al.* 2010; Lang *et al.* 2017). we fix it as a constant 0.5 throughout. This chosen value falls within the empirical range (Lang *et al.* 2017), and using other values within this range does not qualitatively change our simulation results. Similarly, we fix the time cost for failed attacks ($F$) as a constant 0.5 throughout. Looking at the extremes, $F = 0$ means that failed attacks cost the consumer no time at all, whereas $F = 1$ means that failed attacks cost the consumer the full-length of handling time even if the resource item is not captured ($F = 1$). Thus, an intermediate value is likely more realistic. Using other values between 0 and 1 can change the topology of the predicted optimal-foraging food web (i.e., the initial web) all else being equal, but does not change qualitatively the dynamical results.

For the per-capita consumer search rate ($a$, units of area/time or volume/time), we follow the literature (Pawar *et al.* 2012; Tang *et al.* 2014; Rizzuto *et al.* 2018; Ho *et al.* 2021) by adopting a metabolically-constrained mass-scaling rule based on whole-organism consumption. Notably, as our preliminary test shows that using mass-scaling rules of different foraging strategies or foraging dimensions (Pawar *et al.* 2012; Ho *et al.* 2021) does not qualitatively influence our dynamical simulations, we adopt only the mass-scaling rule of 2D active-capture foraging throughout. That is, for species $j$ searching for $i$,

$$a_{ij} = 1.07 \times m_j^{0.21} m_i^{0.21} \sqrt{m_j^{0.42} + m_i^{0.42}} \ , \tag{S1}$$

where $m$ is the species' body mass (in kg, the same hereinafter). We pick active-capture because it represents a more general form of searching, whereas the other two strategies can be seen as the special cases where either the resource or the consumer is immobile. As this mass-scaling rule was derived based mostly on whole-organism predator-prey interactions, by adopting such scaling we focus our modelling exploration on whole-organism feeding and its resultant food webs.

We phenomenologically model the attack success probability of species $j$ attacking $i$ as:

$$A_{ij} = \left( \frac{1}{1 + \left( \log_{10} \left( R_\mathrm{p} \frac{m_j}{m_i} \right) \right)^2} \right)^{0.2} \ , \tag{S2}$$

where $R_\mathrm{p}$ is the preferred resource-consumer body-size (body mass) ratio. This imposes a dome-shaped change in success probability (bounded between 0 and 1) with resource-consumer body-size ratio, with its breadth controlled by the exponent 0.2. The success probability peaks at one when the size ratio of the focal interacting consumer-resource pair equals $R_\mathrm{p}$. Because size ratios tend to be log-normally distributed, the $\log_{10}$ transformation guarantees that the decline is symmetric on either side of this ratio. This model (eqn (S2)) captures the biologically-realistic feature of attack success that the consumer's attack is more likely to succeed when the resource's body size is appropriate, and less likely when the prey is too large or too small (*sensu* Caparroy *et al.* 2000; Vucic-Pestic *et al.* 2010; Pawar 2015). For $R_\mathrm{p}$, as we focused on whole-organism consumption where consumers tend to eat resources smaller than themselves (Brose *et al.* 2006; Pawar *et al.* 2012), we set its value to 0.1

throughout, following previous work (Ho *et al.* 2019, 2021).

According to Pawar *et al.* (2012), the consumer's per-capita handling time for a resource item is positively associated with the body size of the resource while negatively associated with the consumer's. Therefore, we express handling time ($h$) of consumer species $j$ handling resource species $i$ as

$$h_{ij} = h_0 \frac{m_i^{1.25}}{m_j^{1.1}} \ , \tag{S3}$$

where $h_0$ is a scaling constant that reflects the overall magnitude of handling time. Indeed, this captures the nature that for the same consumer, handling larger resource items takes more time than handling small ones, while for the same resource, a larger consumer could handle it much quicker. The 1.1 scaling exponent for the consumer's mass is chosen based on Pawar *et al.* (2012); the 1.25 scaling exponent (slightly larger than one) for the resource's mass is set based on the consideration that the required handling time should increase with the resource's body mass superlinearly, such that handling a 10-kg resource item is more time-costly than handling ten 1-kg items, as the former needs extra time to be dismantled into smaller pieces.

### S2 Food-web structural measures

Here we explain the definition and quantification methods of the food-web structural measures in this study.

The number of top consumers and the mean resource-consumer body-mass ratio are self-explanatory (note that the latter is in $\log_{10}$ scale). Top consumers have been proposed to be influential in determining food-web dynamics (Post *et al.* 2000; Bascompte *et al.* 2005; Michalko & Pekár 2017; Ho *et al.* 2021). As we keep the number of species in synthetic communities to 50 and fix 5 species to be the basal resources, knowing the number of top consumers equals knowing the number of intermediate consumers at the same time. The mean resource-consumer body-mass ratio reflects the overall body-size relationships between consumers and resources within the food web.

Connectance is the the proportion of realised out of all possible (potential) links of the food web, which reflects how connected the web is, or say, the level of trophic link realisation among all species pairings. It is calculated as the number of links in the web divided by the squared species richness. We also quantify the holistic food-web topological measures, nestedness and modularity, of the synthetic food webs. If expressing the food web as a consumer-resource bipartite network, nestedness reflects the degree to which some species' diet comprises a subset of others', whereas modularity reflects the degree of having modules (compartments) whereby many links occur within modules but few occur between them. All these measures have been widely studied for their effects on the system's dynamics (Dunne *et al.* 2002; Krause *et al.* 2003; Kondoh *et al.* 2010).

For quantifying nestedness, we measured the NODF value by using the R package `vegan` (Oksanen *et al.* 2013) with its function `nestednodf` taking the synthetic diet matrices as inputs. For quantifying modularity, we used the R package `igraph` (Csardi & Nepusz 2006) taking the diet matrices of our food webs as inputs, using its function `cluster_walktrap` to derive the module grouping then the function `modularity` to calculate the index reading. Please refer to the manuals of these packages for calculation details.

### S3 Additional figures

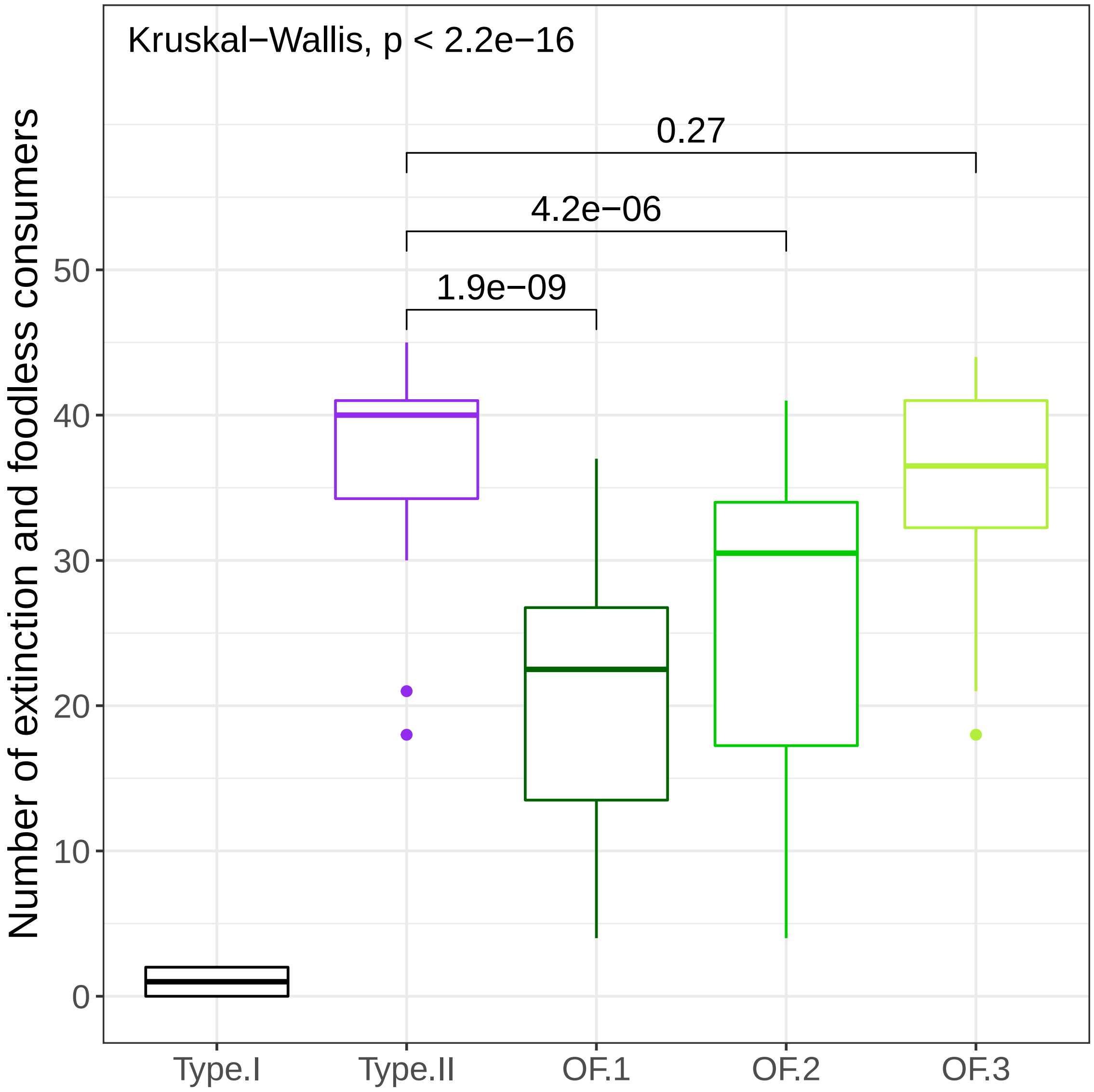


Figure S1: **Number of extinctions and isolated consumers of the five foraging schemes at the end of simulations.** In the Type.I and Type.II schemes (specifically the latter), as consumers have no diet adjustment, some may have lost all their resource species (i.e., biologically cannot maintain) but themselves have not yet fallen below the threshold abundance to be judged as going extinct by the end of our simulations. Here we add up both extinctions and such isolated "foodless" consumers and make Kruskal-Wallis comparisons among the foraging schemes. The global p-value is at the top-left, while we also show pairwise p-values of the comparisons between Type.II and the three optimal-foraging schemes.

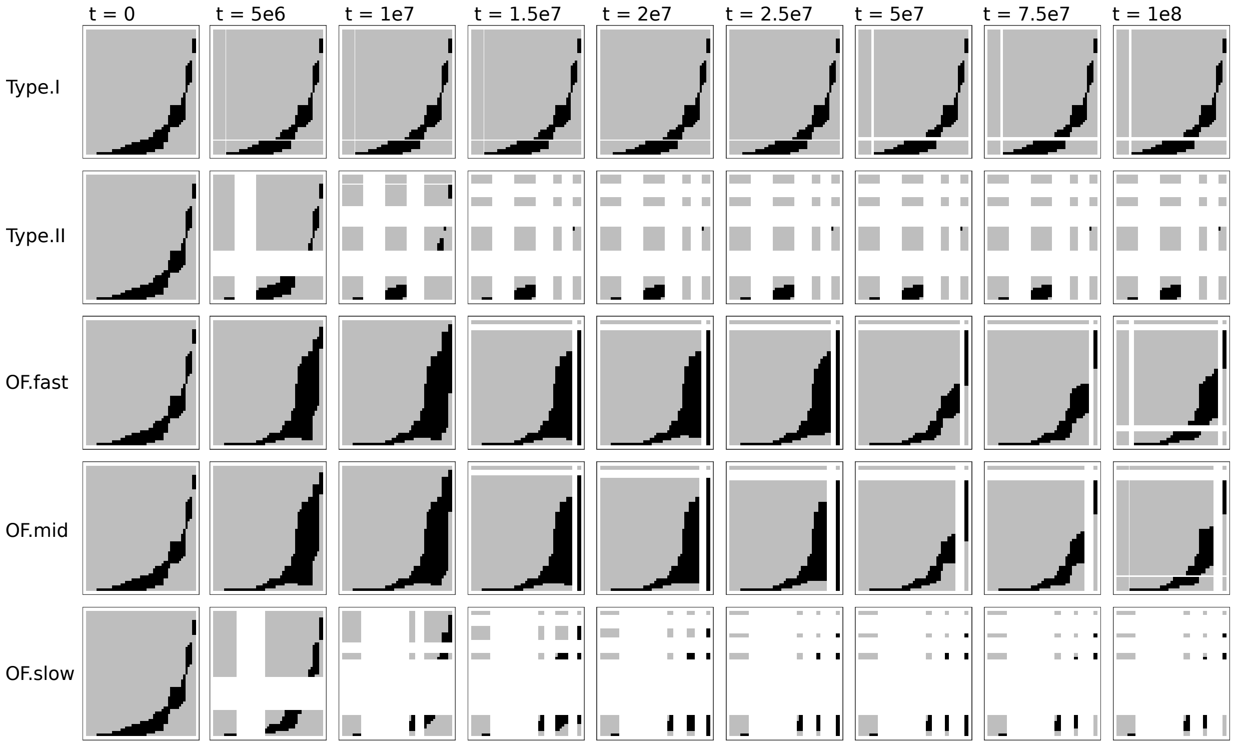


Figure S2: **Diet matrices of a synthetic food web over time of the five diet-adjusting schemes.** Here all matrices are derived from one replicate (i.e., the same community) but the patterns are representative. They are the snapshots of feeding relationships within the food web at the given time points indicated at the top. The black/grey entries respectively indicate realised/non-realised trophic links between consumers and resources (columns and rows, both in small-to-large body-mass order). The white entries indicate that the corresponding species has gone extinct (thus is removed from both the column and row). The left-most five columns have no realised links (not consuming others) as these are fixed to be basal resources. With abundance-dependent trophic interaction strengths (i.e., the Type.II scheme), species rapidly go extinct. Optimal foraging (the OF schemes) lead to changes in the realisation of trophic links over time, and can offset the above rapid extinction, but only when consumers can adjust their diet effectively enough to respond to changing resource abundances (OF.fast and OF.mid, but not OF.slow).

### S4 Additional discussion

In our model, the preferred resource body size for a consumer is set to be a no-larger-than-one ratio to its own body size. Indeed, due to morphological trait-matching constraints of foraging, such as detection or gape-size limitation, the resource-consumer body-size relationships in the real world mostly follow such a pattern (Pawar *et al.* 2012; Eklöf *et al.* 2013; Berlow *et al.* 2008). As a result, the preferred body-size ratio effect (through determining attack success probabilities) embedded in the OF diet choice leads to food webs where consumers mostly eat resources smaller than themselves. Moreover, as we model handling time to increase superlinearly with the resource's body mass, smaller resources are more profitable to the consumer than larger resources with the same extent of deviation from the preferred body-size ratio. In other words, a resource that is ten times smaller than the preferred ratio is more profitable than a resource that is ten times larger than the preferred ratio. Therefore, a consumer's optimal diet, from the narrowest to broadest possibility following resource profitability order, tends to include the small-sized resources before the large-sized ones. Both of these effects contribute to the emergence of food-web nestedness from the OF diet choice mechanism. In our simulation, this happens in the early phase where not too many species have gone extinct, and large consumers broaden their diets to encompass almost all remaining species (Fig. 2 in the main text and Fig. S2).

Conversely, food-web modularity emerges when species abundances allow consumers to have their optimal diets broad enough to include several species around the preferred resource size, but also narrow enough to not include those that deviate much from it. This is especially likely to happen when species body masses in the community exhibit discrete categories and thus the OF diets favour the formation of size-specific trophic interaction modules. In our simulation, this happens in the late phase where some species have gone extinct discontinuously along the body-mass order. Extinctions facilitate the formation of discrete size modules, whereas OF facilitates the realisation of trophic interactions within each module, and together the food-web modularity increases (Fig. 2 in the main text and Fig. S2).
